# Non-invasive assessment of integrated cardiorespiratory network dynamics after physiological stress in humans

**DOI:** 10.1101/2025.03.17.643643

**Authors:** Cecilia Morandotti, Louise Rigny, Thomas B. Williams, Juan I. Badariotti, Matt Miller-Dicks, Amar S. Bhogal, Shachula Han, Jo Corbett, Michael J. Tipton, Joseph T. Costello, Ali R. Mani

## Abstract

**Background:** Physiological variables provide critical insights into the integrated control of the cardiorespiratory system, reflecting the body’s real-time responses to internal and external perturbations. Visualizing the exchange of information between different components of the cardiorespiratory system is beneficial for monitoring individuals in various clinical settings and extreme environments. This study aimed to develop a non-invasive method, using principles of information theory, to visualize the flow of information between physiological variables, with a focus on the integrated cardiorespiratory responses to different physiological stressors.

**Methods:** Heart rate, respiratory rate, minute ventilation, respiratory frequency, tidal volume, capillary oxygen saturation (SpO_2_), end-tidal oxygen, and end-tidal carbon dioxide concentrations were recorded from 22 healthy participants. Transfer entropy, which reflects measures of causal relationships between parallel time-series, was used to compute the flow of information between cardiorespiratory signals into network maps. Network mapping was performed after rest in a control condition and following exposure to the isolated and combined effects of normobaric hypoxia (FIO_2_: 0.12), moderate intensity cycling exercise (100W), and overnight sleep deprivation. For each intervention, 10-minute segments of physiological signals (from minutes 5 to 15 after the commencement of hypoxia and/or exercise) were used for analysis.

**Results:** Each physiological stressor was associated with a distinctive pattern of information flow between physiological variables. Hypoxia led to the engagement of SpO_2_ a hub in the network, facilitating the exchange of information with end-tidal oxygen concentration and heart rate. Sleep deprivation was associated with a shift in the flow of information from SpO_2_ to other nodes, such as respiratory rate, during hypoxia. During exercise, heart rate emerged as the central node for receiving information, while SpO_2_ acted as the primary node disseminating information to other nodes. Increased connectivity within the networks was observed during exercise alone or when combined with other stressors.

**Conclusion:** This non-invasive network mapping technique visualizes the interaction of various cardiorespiratory variables following exposure to physiological stressors. By mapping normative responses, this approach may help identify outliers associated with various disease states.

## Introduction

Network physiology is an interdisciplinary field that investigates the topology of functional interactions between various components of physiological systems. This emerging field has shown promise in mapping the complex interaction among different physiological components in health (Bashan et al., 2012, de Abreu et al., 2023) and has potential to be used for the assessment of patients with critical illnesses and exposure to extreme environments such as hypoxia (Jiang et al., 2021, Oyelade et al., 2024 and Morandotti et al., 2025). Raw physiological data/signals provide insights into the integrated control of the physiological system in both health and disease. The mean or absolute values of single physiological variables, such as heart rate, respiratory rate and peripheral capillary oxygen saturation, are traditionally used to monitor patients in healthcare settings. However, the human body consists of multiple integrated network systems that interact to maintain physiological homeostasis (Bartsch et al., 2015, Garcia-Retortillo and Ivanov, 2024). This dynamic network results in complex fluctuations in physiological signals, which are normally reduced in pathological states when physiological network communication is disrupted (Morandotti et al., 2025). For example, reduced complexity of fluctuation of physiological signals has been shown to predict poor outcome in complex diseases such as sepsis and COVID-19 (Papaioannou et al., 2013, Gheorghita et al., 2022, Alassafi et al., 2024). Most investigators study fluctuations in physiological signals in isolation, using linear and non-linear methods such as approximate and sample entropy (Pincus, 1991, Costello et al., 2020). However, analyzing the interaction between parallel physiological signals provides a more comprehensive view of the control system (Jiang et al., 2021). In fact, visualizing the information exchange between various cardiorespiratory signals would be beneficial for monitoring individuals in different clinical settings (e.g. early warning in critical care) and would provide insights into the integrity of physiological control mechanisms (Morandotti et al., 2025). However, the main limitation for such visualization is the lack of a robust methodology that can be used for network mapping with non-invasively recorded physiological signals.

Information theory studies how information is processed and transmitted, quantifying it by measuring the average amount of uncertainty (or entropy) in a signal. The application of information theory principles (e.g. entropy analysis) offers analytical methods for measuring the flow of information between different physiological variables. A network approach to exploring information transfer between pairs of physiological signals has been applied in the contexts of extreme physiology (Jiang et al., 2021) and critical care (Morandotti et al., 2025). In these studies, the bidirectional flow of information between physiological signals was estimated using Transfer Entropy analysis. Transfer Entropy (TE) determines the causal relationship between variables by measuring the time-directed flow of information from one variable to another (Schreiber, 2000, Wibral et al., 2014). More precisely, TE quantifies the additional information provided by the past segment of one time-series about the future observations of another time-series independent of our knowledge of the past state of the first time-series (Figure 1). For example, in healthy individuals the respiratory and cardiac cycles influence each other, with heart rate (HR) increasing during each inspiration, which is known as respiratory sinus arrhythmia (Yasuma and Hayano 2004, Ben-Tal et al., 2012, Cairo et al., 2023). Thus, there is an exchange of information between the respiratory rate (RR) and HR time-series. Considering two variables, RR and HR, TE quantifies the direct influence of a data segment of RR on the future values of HR. TE computation allows for quantification of the bidirectional relationship between physiological time-series (Feas et al., 2014) and thus the transfer of information can also be measured from HR time-series to RR time-series [TE(HR → RR)] which is mechanistically different from the effect of respiratory cycle on HR [TE(RR → HR)] (Günther et al., 2022). Transfer entropy has been described in various studies as an analytical tool for data-driven causal inference (Barnet et al., 2009, Wibral et al., 2014). Its use can be also extended in network science for mapping the interaction between various nodes (signals) of a physiological system (Figure 1).

**Figure 1.**
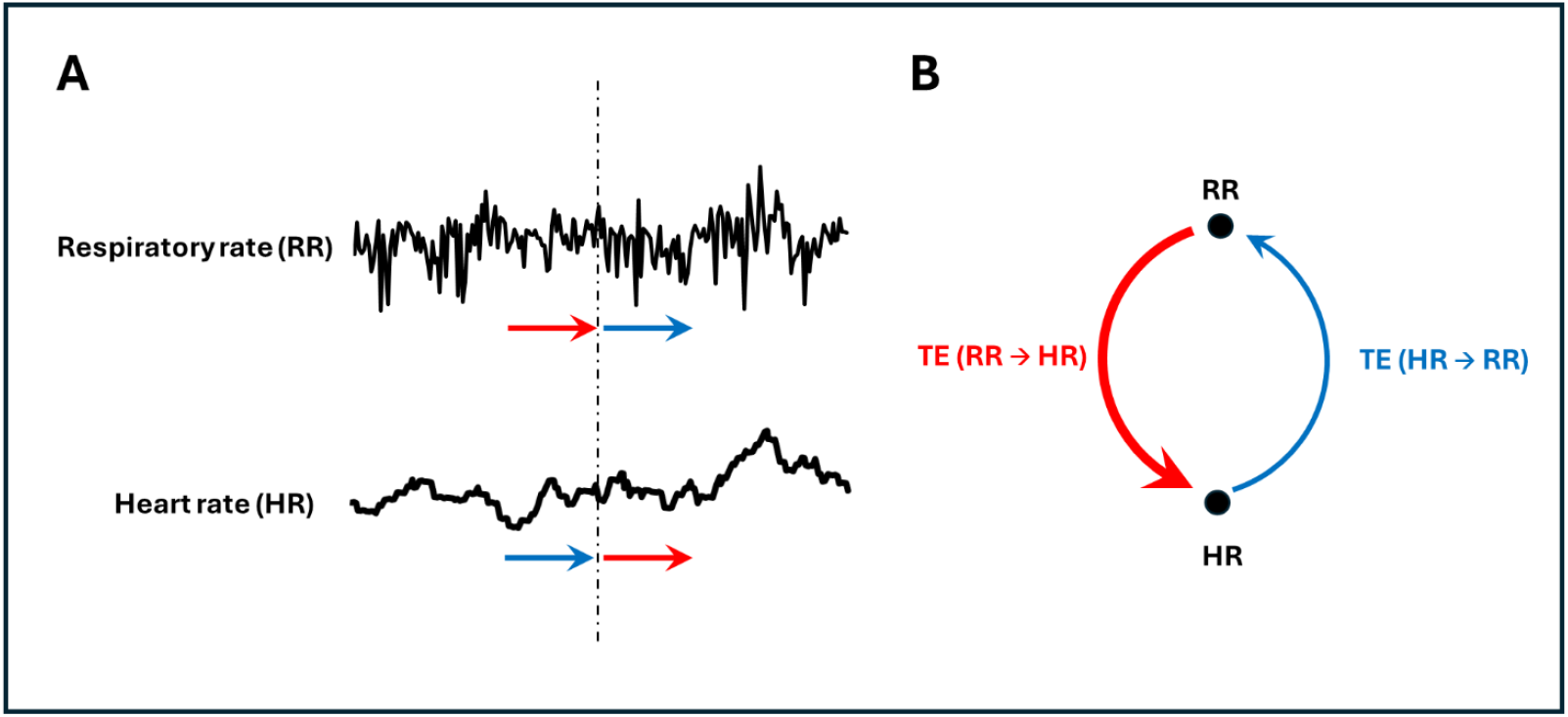
Schematic diagram to explain the concept of transfer entropy (TE). **A**. Example of two parallel physiological time-series, such as respiratory rate (RR) and heart rate (HR), recorded from a participant. **B**. Graphical representation of information transfer between the two physiological signals, where nodes represent the signals and arrows indicate the directional influence of one signal on the other. Integrated physiological mechanisms lead to reciprocal interaction between two parallel physiological signals, e.g., RR and HR. The transfer of information from RR to HR is annotated as TE (RR → HR) quantifies the additional information that the past values of the RR time-series provide about future observations of the HR time-series (represented by red arrows in panel A), independently of information from previous HR states. Likewise, the transfer of information from HR to RR, TE (HR → RR), quantifies the additional information that the past values of the HR time-series provide about future observations of the RR time-series (represented by blue arrows in panel A), independently of information from previous RR states. Such transfer of information can be presented as a network with HR and RR as nodes and TEs as edges (Panel B).

A network mapping method should be able to demonstrate the interaction of multiple physiological components or variables and assess whether the interactions between the nodes are genuine and not due to chance. As no universal network mapping method currently exists, the this exploratory study sought to develop a method to map the dynamic interaction between cardiopulmonary physiological variables, namely, HR, RR, tidal volume (Vt), minute ventilation (Ve), capillary oxygen saturation (SpO_2_), end-tidal oxygen (P_ET_O_2_), and end-tidal carbon dioxide (P_ET_CO_2_). We also tested the hypothesis that physiological stressors such as hypoxia, exercise, and sleep deprivation, both in isolation and combination, altered the dynamics of the physiological network.

## Methods

### Ethics

All participants provided written informed consent with the opportunity to withdraw at any time. A thorough health history questionnaire was completed alongside medical screening prior to participation. The study fully complied with the Declaration of Helsinki, except for registration in a database. All experimental procedures were approved by the Science Faculty Ethics Committee of the University of Portsmouth (project number SHFEC 2021–031 & SHFEC 2018–068).

### Participants

This study was part of a larger project investigating effects of combined stressors on physiological and cognitive function and the experimental design has been described in detail elsewhere (Williams et al., 2024). Data collection was conducted at the Extreme Environments Laboratory, at the University of Portsmouth in the UK. Convenience sampling was employed with healthy, physically fit participants. Data were collected from two experiments conducted at the Extreme Environment Laboratory between 2018-2023. In the first experiment, 12 participants underwent exposure to normobaric hypoxia, moderate intensity exercise, or one night of total sleep-deprivation (either in isolation or combination). In the second experiment, 12 participants underwent exercise or three nights of partial sleep-deprivation (either in isolation or combination). Data from two participants were excluded from the analysis due to incomplete datasets. Consequently, data from a total of 22 (5 females) healthy participants were included in the final analyses. The participants’ characteristics (mean ± SD) were as follows: age: 25.1 ± 3.6 years, height: 177.7 ± 11.7 cm, body mass: 75.4 ± 14.6 kg, body mass index: 23.7 ± 2.4 kg.m^-2^, VO_2_max 44.8 ± 8.0 ml.kg.min^-1^. All participants had a regular sleeping pattern and were free of any cardiovascular, respiratory, cerebrovascular diseases or sleeping disorders, and had not travelled to an altitude higher than 1000 m for at least 1 month prior to commencement of the study, including commercial flights. Participants’ cardiorespiratory parameters, including HR, P_ET_O_2_, P_ET_CO_2_, RR, V_e_ and V_t_ were measured and monitored non-invasively for 45 min using a metabolic cart (Quark CPET, Cosmed, Rome, Italy). SpO_2_ was measured using a pulse oximeter (Nonin 7500, US). Sampling rate for recording physiological variables was one sample per respiratory cycle.

During these trials. physiological signals (i.e., HR, P_ET_O_2_, P_ET_CO_2_, RR SpO_2_, V_e_, V_t_) were measured during different physiological challenges, namely, hypoxia, exercise, and sleep-deprivation, isolated and combined. In brief, the study involved experimental trials, including exercise (cycling at 100W) for 40–minutes under both normoxic (FIO_2_ = 20.93%, sea level) and hypoxic conditions (F_I_O_2_=12.00%, equivalent to ∼4500 m above sea level) following either a full night’s rest, or sleep-deprivation (either overnight sleep-deprivation or three nights of partial sleep-deprivation: For details of the study design, see Williams et al., 2024). Experimental conditions therefore included normoxia control (NC), hypoxia control (HC), exercise control (EC), hypoxia-exercise control (HEC), normoxia sleep-deprived (NS), hypoxia-sleep-deprived (HS), exercise-sleep-deprived (ES), and hypoxia-exercise-sleep-deprived (HES). All participants completed the study. Data from one participant during NS and two participants during ES had noisy HR channel due to detachment of electrodes and were discarded from analyses.

### Data curation

Physiological time-series data with more than 5% missing values were excluded from the study. This exclusion occurred on only 12 occasions (1.26 % of total sample) across different time-series recorded throughout the entire experiment and its interventions. A digital filter was developed to remove any missing values and replace them with the overall average value of the data thread before any calculations. For each intervention, a 10-minute segment of physiological signals (from minutes 5 to 15 after the commencement of hypoxia and/or exercise) were analyzed to calculate TE and network mapping.

### Transfer Entropy calculation and assessment of its significance

Transfer Entropy (TE) was computed by using an open-source MATLAB function (https://www.physionet.org/conTEt/tewp/1.0.0/) based on the Darbellay-Vajda (D-V) adaptive partitioning algorithm as described (Lee et al., 2012). We calculated TE values between all pairs of the following physiological variables; HR, RR, P_ET_O_2_, P_ET_CO_2_, Vt, Ve and SpO_2_, for all TE calculations the time lag was set to 5 respiratory cycles. This choice is based on previous studies that indicated the transfer entropy (TE) between cardiorespiratory signals increases with the time lag and reaches a plateau at a time lag of 15 seconds (Morandotti et al., 2025).

To ensure that the calculated TE values represent genuine directional information transfer between two physiological variables, we applied the Monte Carlo method to distinguish between significant TEs and those values that can be explained by random surrogate data (Figure 2) (Lee et al., 2012, Shirazi et al., 2013). During this process, following the calculation of original TE (A → B), the data points of the source variable’s (A) time-series were randomly shuffled, and the TE was then calculated between the shuffled time series (A) and the authentic time-series of the second variable (B). This process was repeated 100 times to generate the probability distribution of the shuffled time-series TEs. A TE value was considered significant if its value was greater than the 95th percentile of the distribution of TE values derived from shuffled time-series, meaning that the TE value is less likely to be due to randomness. A TE value smaller than the 95th percentile was considered non-significant, or due to chance (with more than 5% uncertainty), hence it was replaced with 0.

**Figure 2.**
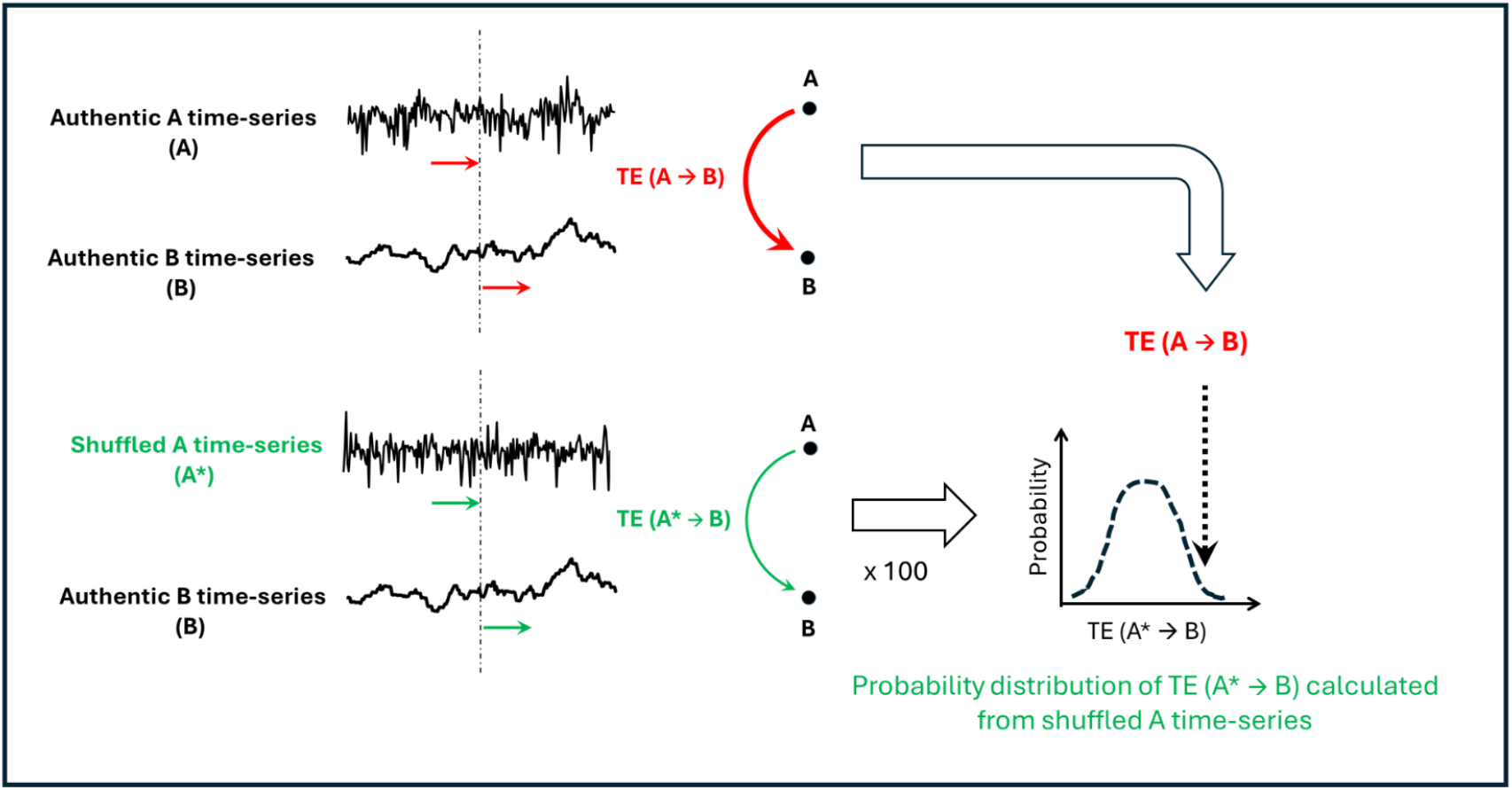
A schematic diagram on the Monte Carlo method for identification of significant information transfer between two parallel physiological time-series, A and B. The source time-series (A) were shuffled, and the Transfer Entropy (TE) was calculated to measure the value of TE between a random time-series and B. This process was repeated 100 times to obtain the distribution of TE calculated from surrogate time-series. The transfer entropy from the original source time-series was deemed significant if it was greater than the 95th percentile of the surrogate results.

### Network mapping and calculation of centrality indices

TEs were calculated between all pairs of physiological variables for each participant in all conditions for which data were available. Medians were then taken of the TE values, and compiled into 8 adjacency matrices, one for each condition/intervention (NC, HC, EC, HEC, NS, HS, ES, and HES). These adjacency matrices were then used to develop network graphs in MATLAB, where the seven physiological variables are represented by nodes, and the directional information transfer is represented by edges, whose thickness is proportional to the TE values. To find the most important nodes that exchange information in each experimental condition, centrality indices were calculated for each node of each participant in each intervention. In network science, indegree (ID) and outdegree (OD) are classic metrics for measuring node “centrality” or “influence” Because information transfer between physiological processes is directional, we applied centrality measures appropriate for directed networks. In these networks, the influence of node A on node B (A → B) is distinct from the influence of node B on node A (B → A); these directions are meaningful and do not cancel each other out. The ID of a node represents directed flow of information coming into that node, indicating how much influence the node receives from other nodes in the network. The OD of a node reflects the flow of information going out from that node, representing how much influence the node sends to other nodes. A MATLAB function was used to calculate ID and OD from the adjacency matrices for all 8 conditions. Network diagrams where node size is proportional to ID and OD values were created using MATLAB.

### Statistical analysis

Physiological variable data are expressed as mean ± SD or median (first-third quartile) based on the distributions of the data. TEs and centrality indices were calculated using MATLAB (MathWorks, MA, USA). Statistical analysis was performed using SPSS Statistics (IBM, Ill, USA). Since the effects of three interventions (hypoxia, exercise, and sleep-deprivation) were assessed, three-way ANOVA was conducted to assess the effect of each condition and their interactions on physiological variables and network indices (e.g. total TE). Some physiological variables and network indices including SpO_2,_ P_ET_O_2_, and V_e_ did not show homogeneity of variances (Levene’s statistic), hence the data was transformed using either log or Box-Cox transformation (for SpO_2_) prior to performing three-way ANOVA. P-values less than 0.05 were considered statistically significant.

## Results

Table 1 shows how the interventions (hypoxia, exercise and sleep-deprivation) affected the mean of physiological variables. Three-way ANOVA showed that hypoxia (P<0.001) and exercise (P<0.001) significantly increased heart rate regardless of sleep condition or combination of the interventions (i.e., no significant interaction between the interventions). Respiratory rate was increased by exercise (P<0.001) regardless of other interventions and their combinations as there was no significant interaction between the interventions (hypoxia, exercise and sleep-deprivation). Three-way ANOVA also indicated that tidal volume (V_t_) was significantly increased by hypoxia (P<0.001) or exercise (P<0.001), whilst there was no significant interaction between hypoxia, exercise and sleep-deprivation. Similarly, hypoxia (P=0.041) had a significant effect on minute ventilation (V_E_) with no interaction between the interventions. Similarly, exercise significantly (P<0.001) effected minute ventilation (V_E_) with no interaction between the interventions. P_ET_O_2_ was significantly reduced by hypoxia (P<0.001) or exercise (P<0.001), regardless of sleep condition or combination of the interventions. P_ET_CO_2_ increased significantly with exercise (P<0.001) regardless of sleep condition, and there was a significant interaction between exercise and hypoxia (P<0.001), indicating that hypoxia affects P_ET_CO_2_ more when combined with exercise compared to hypoxia alone. SpO_2_ was reduced significantly with hypoxia (P<0.001), regardless of sleep condition. There was also a significant interaction between exercise and hypoxia (P<0.001), indicating that hypoxia could reduce SpO_2_ more when combined with exercise, compared to hypoxia alone.

**Table 1.** Mean heart rate (HR), respiratory rate (RR), tidal volume (Vt), minute ventilation (Ve), end-tidal oxygen (P_ET_O_2_), end-tidal carbon dioxide (P_ET_CO_2_) and capillary oxygen saturation (SpO_2_) for all experimental conditions, normoxia control (NC), hypoxia control (HC), exercise control (EC), hypoxia- exercise control (HEC), normoxia sleep-deprived (NS), hypoxia-sleep-deprived (HS), exercise-sleep-deprived (ES), and hypoxia-exercise-sleep-deprived (HES).

|  | NC | HC | EC | HEC | NS | HS | ES | HES |
| --- | --- | --- | --- | --- | --- | --- | --- | --- |
| Sample size | 22 | 12 | 22 | 12 | 22 | 12 | 22 | 12 |
| HR ($\text{min}^{-1}$) | 77.0 ± 12.3 | 86.6 ± 12.0 | 110.8 ± 12.4 | 122.5 ± 14.0 | 79.8 ± 9.4 | 87.1 ± 9.2 | 115.3 ± 11.5 | 126.6 ± 9.9 |
| RR ($\text{min}^{-1}$) | 17.8 ± 3.0 | 16.8 ± 3.5 | 26.5 ± 3.2 | 25.1 ± 4.2 | 18.5 ± 2.4 | 16.8 ± 3.7 | 26.6 ± 3.8 | 26.2 ± 5.0 |
| $V_t$ (L) | 0.86 ± 0.17 | 1.02 ± 0.20 | 1.50 ± 0.23 | 1.74 ± 0.18 | 0.84 ± 0.20 | 1.04 ± 0.22 | 1.47 ± 0.25 | 1.56 ± 0.18 |
| $V_e$ (L) | 14.0 ± 2.3 | 15.4 ± 2.5 | 38.9 ± 6.7 | 41.7 ± 5.8 | 14.0 ± 2.5 | 15.4 ± 2.9 | 36.7 ± 4.9 | 38.5 ± 8.1 |
| $P_{ET}O_2$ (mmHg) | 110.4 ± 3.7 | 52.8 ± 2.9 | 105.0 ± 3.0 | 48.1 ± 1.4 | 110.6 ± 5.0 | 52.6 ± 2.5 | 105.4 ± 3.2 | 50.4 ± 3.3 |
| $P_{ET}CO_2$ (mmHg) | 34.1 ± 2.6 | 33.5 ± 2.5 | 40.2 ± 2.8 | 36.5 ± 2.2 | 34.2 ± 3.5 | 34.0 ± 2.6 | 40.6 ± 2.9 | 36.6 ± 2.1 |
| $SpO_2$ (%) | 99.6 ± 0.6 | 87.9 ± 2.9 | 99.5 ± 0.7 | 78.2 ± 3.4 | 99.8 ± 0.4 | 87.0 ± 2.2 | 99.6 ± 0.4 | 78.8 ± 3.0 |

Figure 3 presents network diagrams illustrating the flow of information between physiological signals under different experimental conditions. In these diagrams, the edges represent the median of significant information transfer between physiological variables (nodes). The diameter of each node corresponds to its indegree, reflecting the total amount of information received by that node. Network maps where node diameters indicate outdegree, representing the total amount of information sent out by each node, are provided in Appendix 1.

**Figure 3:**
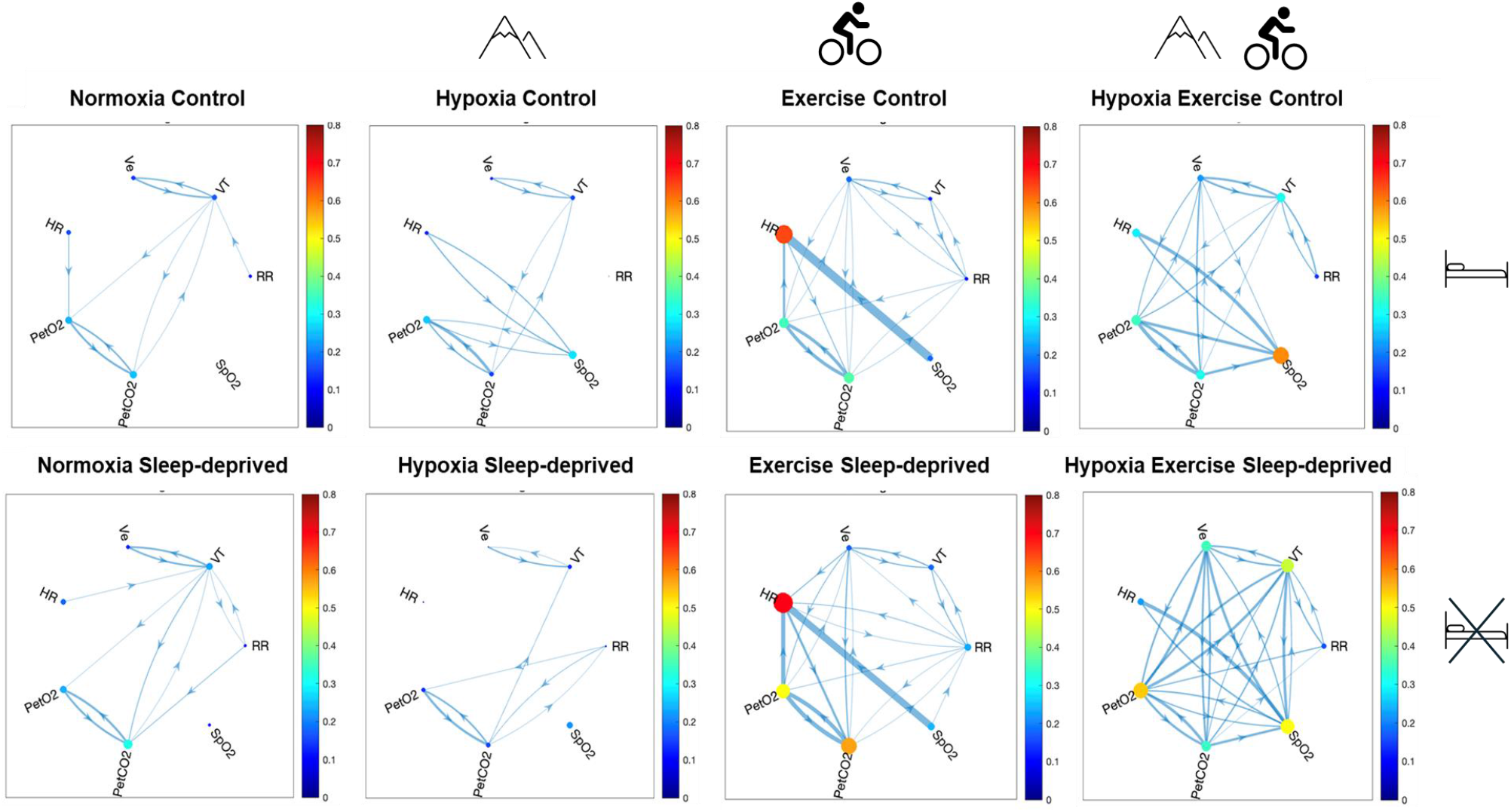
Network mapping based on flow of information transfer between seven physiological variables heart rate (HR), respiratory rate (RR), tidal volume (Vt), minute ventilation (Ve), end-tidal oxygen (P_ET_O_2_), end-tidal carbon dioxide (P_ET_CO_2_) and capillary oxygen saturation (SpO_2_) in different experimental conditions, normoxia control (NC), hypoxia control (HC), exercise control (EC), hypoxia- exercise control (HEC), normoxia sleep-deprived (NS), hypoxia-sleep-deprived (HS), exercise-sleep-deprived (ES), and hypoxia-exercise-sleep-deprived (HES). The thickness of the edges is proportional to the median transfer entropy (TE) value, and only significant TE values are represented by the edges. The size of each node is proportional to its ***indegree*** centrality.

Network mapping showed that in health in the normoxic non exercising control condition (NC), there were three prominent bidirectional transfer of information between P_ET_O_2_ ⇌P_ET_CO_2_, Ve ⇌Vt, and Vt ⇌P_ET_CO_2_. Significant unidirectional transfer of information was also observed between HR → P_ET_O_2_, RR→ Vt, and Vt→P_ET_CO_2_. This is reflected by the ID and OD values of the network, where nodes which receive the most information were P_ET_O_2_ and P_ET_CO_2_ (Figure 3), whereas nodes with the highest output of information were P_ET_O_2_, P_ET_CO_2_ and HR (Appendix 1).

In the presence of physiological interventions, we observed changes in the interaction patterns and the flow of information between most nodes in the network. In HC conditions, additional bidirectional information transfer appeared between HR and SpO_2_ compared to NC. Transfer of information from P_ET_CO_2_ to P_ET_O_2_ also became more prominent in HC. There was also an increase in the amount of information received by SpO_2_ from HR and P_ET_O_2_ reflected in relatively high ID of SpO_2_ node during HC (Appendix 1). P_ET_O_2_ and P_ET_CO_2_ remained prominent nodes in terms of centrality indices (ID and OD) during NC.

During exercise the flow of information transfer increased as shown in Figure 3 and 4. The most prominent increase in information transfer is from SpO_2_ → HR, and these two nodes become the most prominent of the network, with HR and SpO_2_ having the highest ID and OD respectively. Compared to NC and HC, there is also some increase in information transfer between P_ET_O_2_ and P_ET_CO_2_, as well as their ID and OD values which become more prominent. There was also some transfer of information to and from RR with other nodes, which was not observed in HC and NC. When hypoxia and exercise were combined (HEC), there was an even greater number of links between the nodes. The node with the most input of information was SpO_2_, whereas the nodes with the most output of information were P_ET_CO_2_, P_ET_CO_2_, P_ET_O_2_ and SpO_2_. During HEC, HR received less information than in EC conditions as shown in Figure 3.

**Figure 4:**
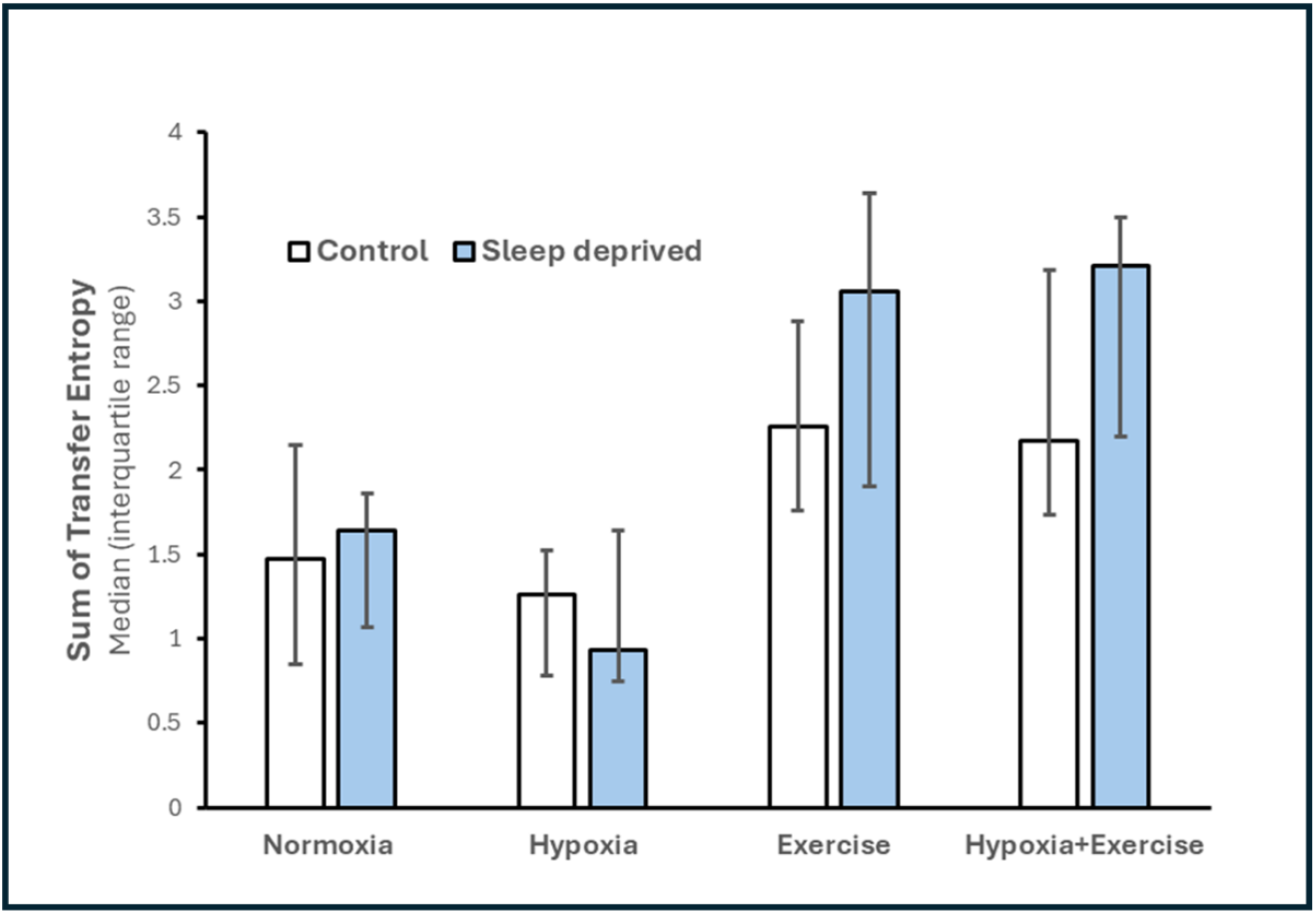
Bar graph showing the median of total sum of transfer entropy during different interventions. Error bars represent the interquartile range. Blue represents control conditions (no sleep-deprivation), and yellow represents sleep-deprivation conditions.

Although three-way ANOVA showed that sleep-deprivation did not significantly affect the mean of any physiological variable, there were differences in the sleep-deprivation network diagrams compared to those of control conditions (Figure 3). In NS, there was prominent flow of information between P_ET_O_2_ and P_ET_CO_2,_ with P_ET_CO_2_ acting as a node with the highest indegree. When hypoxia and sleep-deprivation were combined (HS), there was a shift in flow of information from SpO_2_ (as a central node in NC) to other nodes such as RR. In this condition (HS), the nodes which received the highest amount of information were Vt, P_ET_O_2,_ and P_ET_CO_2_, whilst the node which sent out the most information was RR (Appendix 1). When exercise and sleep-deprivation were combined, the network showed similarities with the EC condition, with the most prominent transfer of information was from SpO_2_ → HR. Consequently, the node with the highest input of information was HR, and the nodes which release the highest amount of information were P_ET_O_2_, SpO_2_ and HR (Appendix 1). In hypoxia-exercise-sleep-deprivation conditions, more links between the nodes appear, with the information transfer evenly spread between all nodes. P_ET_O_2_ was the node which received the most information, with SpO_2_ and Vt receiving the second and third highest amount. Ve, VT, P_ET_O_2_ and P_ET_CO_2_ are the nodes with the highest output of information.

Similarly, as to control conditions (HEC), HR also showed a lower input and output of information compared to exercise without hypoxia (ES and EC). To confirm this, we used three-way ANOVA to assess the effect of interventions on ID centrality measure of HR. The results indicated that ID of HR increased significantly with exercise (P<0.001) and there was a significant interaction between exercise and hypoxia (P<0.001), indicating that exercise alone affects centrality of HR differently than exercise combined with hypoxia.

Figure 4 shows the total transfer entropy values during the three physiological challenges. A three-way ANOVA revealed that exercise is the only intervention that significantly increases the sum of information transfer between the physiological variables recorded in this study (P<0.001), regardless of other interventions, as there were no statistically significant interactions between the interventions on total TE.

## Discussion

In this study, we demonstrated that physiological time-series data recorded non-invasively can be used to map the interactions between different components of the cardiorespiratory system at rest, during exercise and after both hypoxia and sleep deprivation. Healthy young adults exposed to hypoxia, exercise, or sleep-deprivation (both in isolation and in combination), and the flow of information between physiological variables was quantified using an entropy-based algorithm. These interventions are common stressors with efficient adaptive mechanisms that have been investigated using reductionistic methods over the last century (Haldane and Priestley 1905, López-López et al., 1997, McDermott et al., 2006, Millet et al., 2012). While reductionistic methods have been extremely fruitful in elucidating the fundamental physiological mechanisms involved in response to physiological and pathological challenges, they require the isolation/manipulation of physiological subsystems, which limits non-invasive, comprehensive assessment of integrative control mechanisms. The present study demonstrates that the application of information theory principles enables the mapping of interactions within the cardiorespiratory system using non-invasively recorded physiological signals. An advantage of this approach is that it can reveal bidirectional interactions that exist between different components of the cardiorespiratory system and can also distinguish statistically significant information flow from noisy interactions due to stochastic fluctuations of the variables. For the first time, we have demonstrated that each physiological stressor is associated with a distinctive pattern of information flow between physiological variables. For example, hypoxia alone led to the engagement of SpO_2_ (representing peripheral oxygenation) as a hub in the network, exchanging information with P_ET_O_2_ and HR. Conversely, during exercise, HR became the central node for receiving information, while SpO_2_ acted as the main node that sent out information to other nodes. We also observed a different pattern of information flow when hypoxia and exercise were combined. In the HEC condition, SpO_2_ became the main node that received and disseminated information to other nodes. These novel findings extend our existing knowledge about integrated physiological control and provide novel mechanistic insights into the dynamics of integrated cardiorespiratory control in health and disease.

The method described in this study enables us to qualitatively observe the patterns of information flow between physiological signals and identify the most central physiological variable in response to stress, at the level of both the individual and the group. For example, our results indicate that sleep-deprivation may alter the physiological response when an individual is hypoxic. In sleep-deprived individuals, hypoxia led to respiratory rate engagement with P_ET_CO_2_ and Vt, in contrast to the control condition, where hypoxia was associated with the exchange of information with SpO_2_ (Figure 3). While the exact mechanism(s) responsible for these findings are yet to be elucidated, network mapping with TE can reveal bidirectional causal relationships between cardiopulmonary variables, extending the findings of the pattern analysis of SpO_2_ fluctuations during environmental or pathological challenges (Jiang et al., 2021, Gheorghita et al., 2022). Moreover, this method holds potential for the visualization of organ system control in health and disease, and for the monitoring of patients in acute clinical settings (e.g., intensive care units), where multivariate physiological recordings are available. Continuous physiological signals, such as HR, RR, and SpO_2_, are often available in intensive care units. A simplified version of transfer entropy (TE)-based network mapping has recently been reported in critically ill patients with sepsis (Morandotti et al., 2025). This report from our team showed that reductions in TE (HR → RR) and TE (RR → HR) were associated with an increased risk of 48-hour deterioration and mortality, independent of age, mechanical ventilation status, disease severity, and comorbidities (Morandotti et al., 2025). However, this report only considered routine physiological signals and did not use the Monte Carlo method to differentiate between significant TEs and values that could be explained by random fluctuations. The method presented in this study has the potential to be applied to the assessment of critically ill patients, facilitating the development of early-warning physiological biomarkers for deterioration, prognostication or response to therapy. Physiological network mapping using a 10-minute window of physiological signals enables rapid assessment of individual patients. This contrasts with commonly used assessment methods, which typically rely on daily laboratory tests (e.g., SOFA score). Such methods could help clinicians accurately monitor the cardiorespiratory control system to avoid the liberal over-administration of oxygen during ventilatory support, a known complication in current clinical practice (Chu et al., 2018, Gheorghita et al., 2022).

Integrated cardiorespiratory control relies on the communication between central and peripheral chemo- and baroreceptors to maintain a precise balance between oxygen supply and demand. When oxygen levels drop, the respiratory centers in the brainstem adjust their firing patterns to modify breathing rate and volume (Jubran & Tobin, 2000). These physiological responses depend on effective information transfer among various components involved in maintaining tissue oxygenation, including the brainstem, autonomic and cardiovascular centers, cardiac pacemaker, vasculature, airways, and respiratory muscles. Recent evidence suggests that the cardiorespiratory control system’s response to environmental or pathological challenges can be measured by analyzing the pattern of physiological signals fluctuations (Raoufy et al., 2016, Raoufy et al., 2017, Satti et al., 2019, Jiang et al., 2021, Borovkova et al., 2022, Cairo et al., 2023, Romanchuk 2023). Our current findings extend our current understanding of the information transfer between different components of the cardiorespiratory control system, even in the absence of significant changes in the average values of physiological signals (e.g., SpO_2_ during exercise) that are typically used for interpretation of data in physiology and biomedical disciplines.

The application of centrality measures in network science allows for a more comprehensive analysis of network dynamics, building on the research conducted by Jiang and colleagues, where differential network maps were used to hypothesize which variable acted as the physiological ‘hub’ of the mapped networks (Jiang et al., 2021). In this earlier study by our group (Jiang et al., 2021), we used the concepts of indegree (ID) and outdegree (OD) in directed networks to identify the most central nodes in the functional physiological network. SpO_2_ OD generally increased during hypoxic challenges, except when combined with sleep-deprivation (Figure 3). This finding suggests that fluctuations in peripheral oxygen levels are not random but rather reflect the transfer of information from other variables to functionally adapt to hypoxia. During exercise, the SpO_2_ ID ranked low, with heart rate emerging as the most important node for receiving information from other nodes during the EC and ES interventions. This aligns well with exercise physiology, where an increase in cardiac output, primarily attributed to a rise in heart rate, is a key adaptive mechanism. Interestingly, SpO_2_, along with P_ET_O_2_, appeared as an important node with high OD, sending out information that affects heart rate during exercise. The dynamic changes observed when hypoxia and exercise are combined reflect the complexity of integrative physiological control under extreme conditions. The reason why SpO_2_ centrality does not rank highly in the HS group remains unclear. Sleep is crucial for maintaining autonomic, neural, and thermoregulatory function (Albrecht 2012, Morf and Schibler 2013, Kumar et al., 2024). It is well established that sleep-deprivation impairs the synchrony between various physiological subsystems (Healy et al., 2021) and is associated with a reduction in high-frequency HR variability, as well as vascular and metabolic dysfunction (Zhong et al., 2005, Sauvet et al., 2010, Papadakis et al., 2022). These changes reflect alterations in autonomic and metabolic control, which are linked to adverse cardiovascular complications (Krittanawong et al., 2020Azboy and Kaygisiz, 2009). This disruption in normal system functioning may alter communication between physiological functions. Further research is needed to determine whether this was an outlier or if it represents a significant functional change in the network.

In this study, total network TE was calculated for each intervention by summing the TE values between physiological indices. A significant increase in the total sum of TEs was observed in the exercise groups (EC, ES, HEC, and HES) compared to their controls (Figure 4). No differences were observed with hypoxia or sleep-deprivation at rest. The increase in total TE during exercise indicates that exercise requires a substantial increase in adaptation with the cardiorespiratory system. Thus, while the total information content does not change during an acute normobaric hypoxic challenge or sleep-deprivation alone, the transmission between nodes does, suggesting the presence of distinctive compensatory mechanism for each challenge. This hypothesis is further supported by the computed in-degree and out-degree network heatmaps (Figure 3 and Appendix 1), which provide additional insights into the ‘receivers’ and ‘senders’ of information within the network. This additional insight is important as it cannot be achieved by other entropy-based measures such as sample entropy (Bhogal and Mani, 2017, Costello et al., 2020) or other information-based indices such as mutual information (Tan et al., 2020).

The physiological network maps revealed notable visual changes in bidirectional links between physiological signals in the network graphs during interventions, suggesting that the cardiorespiratory network can adapt to different stressors. Previous studies have shown that the functional connectivity of organ systems is crucial for the survival of critically ill patients (Asada et al., 2016, Morandotti et al., 2025). Although our current observations are limited to young healthy individuals, it would be valuable to extend the visualization of the system to disease states where integrated cardiorespiratory mechanisms contribute to patient survival, such as in COPD, sleep apnea, sepsis and acute mountain sickness.

Network dysfunction during disease has been reported in other organ systems. For example, the use of a parenclitic network approach predicted survival outcomes in chronic liver disease patients and provided insights into liver network disruption (Zhang et al., 2022, Oyelade et al., 2023, Oyelade et al., 2024).

Therefore, it is reasonable to suggest that the development of functional network mapping methods could be used to visualize and gain further insights into other organ network systems in both health and disease. Such a network approach can be used for mapping normative responses and may help identify outliers associated with various disease states or susceptibility to potentially catastrophic illnesses (Moss et al., 2016).

### Study limitations

The study is limited to young, healthy, and physically active individuals, who were screened for ethical reasons. As such, our findings may not accurately reflect those of a wider population. Future studies are needed to confirm these findings and to address the study’s limitations by including older individuals and those with common comorbidities. This could be extended to other physiological stressors and clinical settings to assess its utility in monitoring or predicting disease states. Another limitation of this study is that the sampling rate for recording physiological variables was one sample per respiratory cycle. As a result, information transfer at higher frequencies (e.g., information flow leading to high-frequency heart rate fluctuations) could not be assessed. However, if applied to high frequency time-series, the method allows users to assess information transfer at higher frequency. Paired or repeated measures statistical analysis was not possible in this study because data on the hypoxic challenge were not available for all participants (only 12 participants underwent the hypoxic challenge). Future studies could involve paired comparisons with a larger (e.g. with power calculation), more gender- and age-balanced sample.

## Conclusion

We have demonstrated that transfer entropy analysis of parallel physiological time-series offers a method for monitoring integrated cardiorespiratory control and investigating multivariate data-driven causal relationships, where reductionist approaches were previously relied upon. This non-invasive network mapping technique visualizes the interaction of various cardiorespiratory variables after exposure to physiological stressors, and by mapping the normative responses may help identify outliers associated with various disease states.

## Acknowledgements

We would like to express our gratitude to all the participants involved in contributing their time to support this study.

## Data availability

Data will be made available upon reasonable request. Author contribution: J.T.C., T.B.W. and A.R.M. conceived and designed research; T.B.W., and J.I.B collected data; C.M., L.R., A.S.B., S.H. and A.R.M. analyzed data; C.M., L.R., J.C., M.J.T. J.T.C. and A.R.M. interpreted results of experiments; C.M., L.R., and A.R.M. prepared figures; C.M., L.R. and A.R.M. drafted manuscript; S.H., A.S.B, J.T.C, M.J.T., M.M.D and A.R.M edited and revised manuscript; all authors approved final version of manuscript.

## Appendix 1

Network mapping based on flow of information transfer between seven physiological variables heart rate (HR), respiratory rate (RR), tidal volume (Vt), minute ventilation (Ve), end-tidal oxygen (P_ET_O_2_), end-tidal carbon dioxide (P_ET_CO_2_) and capillary oxygen saturation (SpO_2_) in different experimental conditions, normoxia control (NC), hypoxia control (HC), exercise control (EC), hypoxia- exercise control (HEC), normoxia sleep-deprived (NS), hypoxia-sleep-deprived (HS), exercise-sleep-deprived (ES), and hypoxia-exercise-sleep-deprived (HES). The thickness of the edges is proportional to the median transfer entropy (TE) value, and only significant TE values are represented by the edges. The size of each node is proportional to its ***outdegree*** centrality.

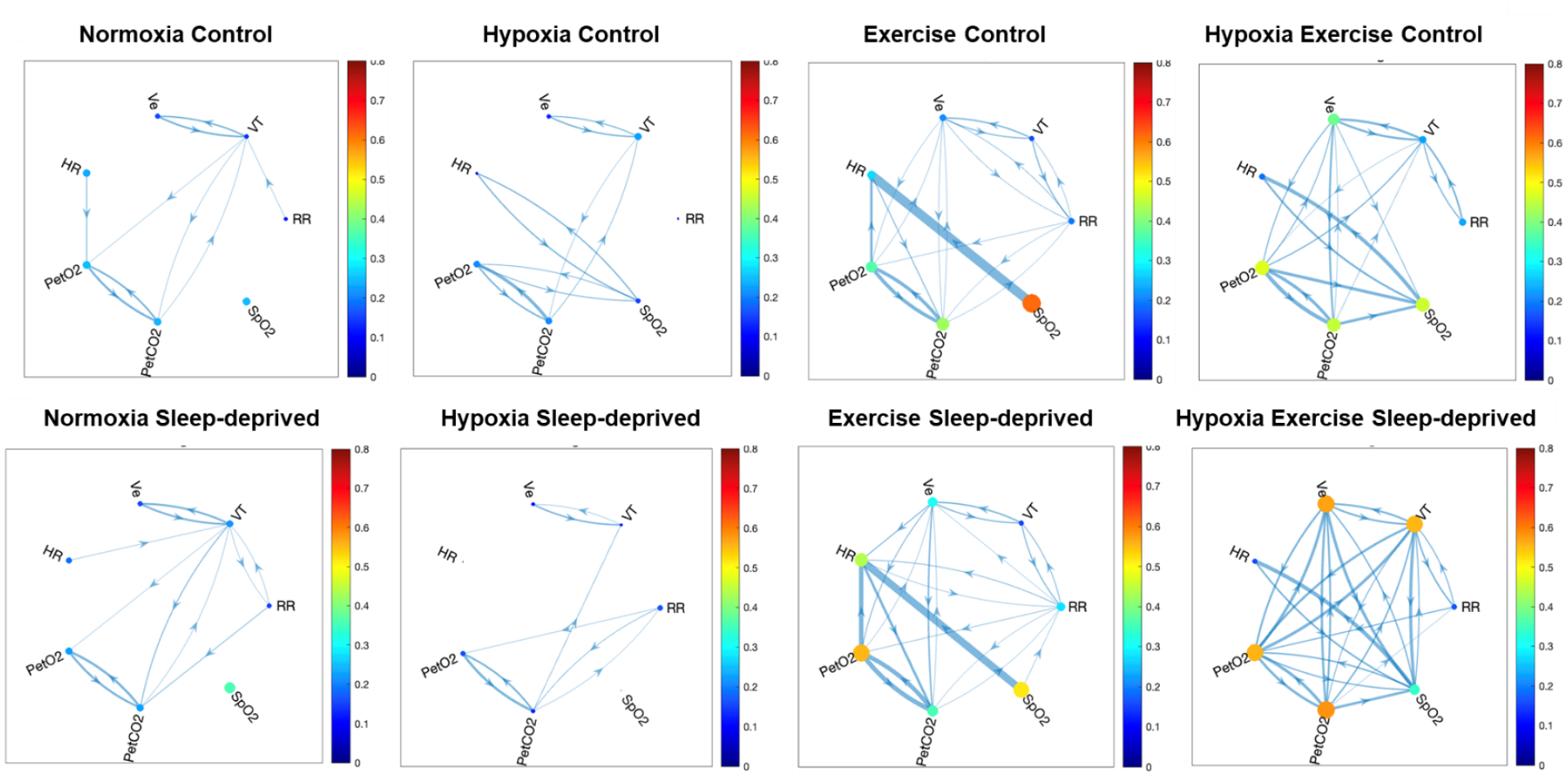

